# Off-target effects as confounders of Cas13d-based lncRNA screens

**DOI:** 10.1101/2025.11.27.690751

**Authors:** Juan J. Montero, Riccardo Trozzo, Roland Rad

## Abstract

Despite their recognized role in biology, a majority of the ∼100,000 lncRNA genes remain functionally uncharacterized. In a recent study (*Liang WW et al*., *Transcriptome-scale RNA-targeting CRISPR screens reveal essential lncRNAs in human cells, Cell, 2024)*, Liang et al. utilized the RNA nuclease Cas13d to perturb ∼6,200 lncRNAs in fitness screens across five cell lines - thereby identifying 778 lncRNAs with broad or context-specific essentiality. However, previous screens reported a lower proportion of essential lncRNAs. To investigate this discrepancy, we re-analysed Liang et al.’s data and found that 68.1% of gRNAs causing fitness defects have off-targets in essential protein-coding genes. This caused numerous false-positive hits, particularly among lncRNAs classified as broadly essential. Off-target effects also compromise the study’s validation efforts, including experiments combining single-cell transcriptomics and lncRNA-perturbations, which confirm the downregulation of off-target protein-coding genes identified in our analyses. The large number of false-positive hits reported by Liang et al. undermines the study’s biological conclusions and endangers future research building on these data, if not considered.

## Introduction

Long non-coding RNAs (lncRNAs) are a heterogeneous class of transcripts that lack an open reading frame and do not encode proteins^1,2^. In humans, lncRNA genes constitute a large part of the genome and outnumber protein-coding genes by approximately 4.5 times^3^. Despite their recognized importance in various biological processes, only a small fraction of lncRNAs has been functionally characterized. However, new high-throughput screening tools are promising to advance the field by facilitating systematic functional interrogation of the lncRNAome.

Several approaches for lncRNA perturbation have been developed, including Cas9-mediated lncRNA deletion^4^ or splice acceptor/donor site editing^5^, CRISPRi transcriptional repression^6^ or targeting of lncRNA transcripts using RNAi. However, each of these approaches has limitations. Cas9 and CRISPRi can cause collateral perturbation of overlapping coding and regulatory DNA sequences. This is not an issue for RNAi, which directly targets RNA molecules. However, RNAi is limited by its primary activity in the cytoplasm^7^, whereas most lncRNAs exert their effects in the nucleus^8^.

We previously addressed this challenge by a screening approach based on the RfxCas13d RNA nuclease, thereby targeting 158,816 lncRNA transcripts encoded by 24,171 lncRNA genes^9^. RfxCas13d directly targets lncRNA transcripts in the nucleus, thus overcoming several limitations of other perturbation methods. A similar approach has been later reported by Liang et al., who developed a screening library targeting ∼6,200 lncRNAs^10^, thereby identifying 778 lncRNAs with broad or context-specific essentiality. However, previous screens using Cas13d or orthogonal approaches reported a substantially lower proportion of essential lncRNAs^6,9^. For example, despite the much larger size of our Cas13d-based screens^9^, we found ∼3.8 times fewer core-essential lncRNAs than Liang et al.

To investigate the causes of this discrepancy, we reanalyzed Liang et al.’s data and found large numbers of false-positive hits due to off-target effects. We show that on average 68.1% (63.3%-76.0%) of gRNAs producing fitness phenotypes in individual screens display off-target effects in essential protein-coding genes. This issue severely compromises the reported list of essential lncRNAs and the study’s biological conclusions.

## Results

### Off-target gRNAs account for the majority of phenotypic gRNAs in the screens

Liang et al. conducted pooled RfxCas13d-based fitness screens to identify lncRNAs involved in cell survival or proliferation. To this end, they developed a lentiviral gRNA library targeting ∼6,200 lncRNAs, which was used to map fitness phenotypes in five RfxCas13d-engineered cell lines. In these experiments, lncRNAs affecting cell viability can be identified by assessing gRNA frequencies at the start and end of the experiment followed by calculation of log-fold change values (LFC) and statistical analyses. The study reports the identification of 778 lncRNAs affecting cellular fitness, hereafter referred to as lncRNA “hits” or lncRNAs identified as “essential”. These lncRNA hits have been categorized by Liang et al. based on their recurrence as cell-type-specific (dropout in one cell line, 61.3%), partially shared (dropout in two to four cell lines, 32.8%), or shared (dropout in all cell lines, 5.9%).

When examining the list of significant hits (Table S2 in Liang et al.), we first noticed that large numbers of non-expressed lncRNAs were detected as hits in each cell line (Figure 1a, Table S1). In total, we found 572 lncRNA hits that are not expressed in any of the cell lines in which they were detected as essential (Figure 1b). These lncRNA hits are not being mentioned by Liang et al. in their main text. A subset of these lncRNAs (n=154) appears as hits across several or even all cell lines, despite not being expressed (Figure 1b). These findings are alarming, as non-existent lncRNA transcripts can neither be essential to a cell nor can they be targeted by RfxCas13d. Hence, hits affecting non-expressed lncRNAs have to be considered false-positive.

**Figure 1:**
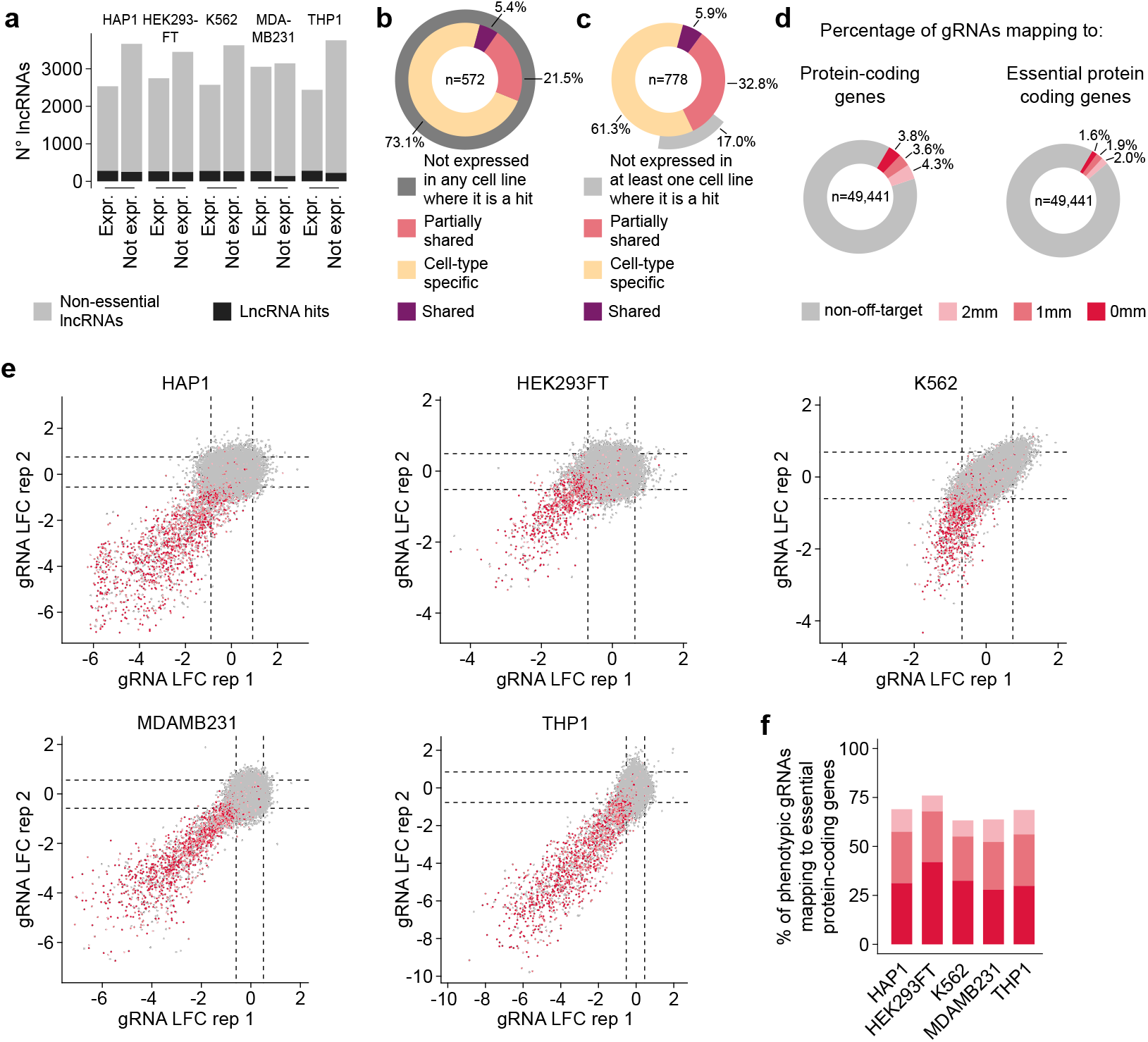
A large part of phenotypic gRNAs in Liang et al.’s screens have off-target effects in essential protein-coding genes. **a**, Number of significant lncRNA vulnerabilities (lncRNA hits, black) and non-essential lncRNAs (grey) identified in each cell line screened by Liang et al. Within each cell line, lncRNAs are further categorized depending on their expression (expressed or not expressed). **b**, Number of lncRNA hits that are not expressed in any of the cell line(s), in which they were identified as essential by Liang et al. Data from Liang et al.’s Table S2.These non-expressed lncRNA hits are further categorized based on their recurrence as cell-type-specific (dropout in one cell line), partially shared (dropout in two to four cell lines), or shared (dropout in all cell lines). **c**, Categorization of the 778 lncRNA hits reported by Liang et al., based on their recurrence across cell lines (cell-type-specific, partially shared, or shared) as described in Liang et al.’s study. The outer circle (grey) indicates, for each category the number of lncRNAs for which expression is absent in at least one cell line in which they were identified as a significant hit. **d**, Percentage of gRNAs in Liang et al.’s library mapping to exons of at least one protein-coding gene (left panel) or to at least one essential protein-coding gene (right panel). Transcript sequences are from GENCODE v38, essential genes from DepMap release 24Q2. The red color scheme indicates gRNAs with a perfect match to their off-target genes (0mm) and gRNAs aligning with one (1mm) or two (2mm) mismatches. **e**, Replicate correlation of log-fold change (LFC) values for gRNAs targeting lncRNAs in Liang et al.’s screens. Dashed lines indicate 95% confidence intervals, computed using the distribution of non-targeting gRNAs (analogous to Liang et al., Figure 1B-C and Figure S1F-J). gRNAs with LFC values below this interval in both replicates are considered “phenotypic” (gRNAs creating robust fitness defects). Off-targets in essential protein-coding genes are indicated for each gRNA by the respective dot color. The red color shades highlight 0mm, 1mm and 2mm as in (d). **f**, Percentage of “phenotypic” gRNAs with off-targets in at least one essential protein-coding gene. Color scheme highlighting 0mm, 1mm and 2mm as in (d).

Examining the 778 essential lncRNAs reported by Liang et al, we found a similar issue for 17% of hits (n=132) (Figure 1c). These lncRNAs are hits in multiple cell lines but lack expression in at least one of them. Thus, a total of 704 (572+132) lncRNA hits are likely artefactual (Figure 1b, 1c). Moreover, since false-positive rates are expected to be similar across the entire screen, we envisaged that a part of the expressed hits might also be false-positive.

We hypothesized that false-positive hits could stem from gRNAs with off-target effects in protein-coding genes, which may cause fitness defects, even if the intended lncRNA target is not expressed. We therefore queried potential off-targets for all gRNAs in the library using the criteria outlined by Liang et al and published earlier by the same group^10,11^. Specifically, gRNAs were mapped to exons of all human protein-coding transcripts, followed by exclusion of alignments to the forward-reference strand. gRNAs with up to two mismatches to protein-coding gene sequence were retained and considered gRNAs suffering from off-target effects. We found that 11.7% (n=5,757) of gRNAs in the library had off-targets in protein-coding genes, with approximately one third of them showing a perfect match (Figure 1d, Table S2). This suggests issues with off-target filtration during library design. Most problematic are off-targets in common essential protein-coding genes, as their perturbation causes cellular fitness defects^12^. This problem affects 5.5% (n=2,726) of all gRNAs in the library (Figure 1d).

To examine the potential implications of this observation, we assigned off-target information to Liang et al’s screening results. This revealed that on average 68.1% of gRNAs producing fitness defects (referred to as “phenotypic gRNAs”), have off-targets in essential protein-coding genes (Figure 1e,f). Thus, inaccurate filtration of off-target gRNAs creates a noteworthy confounding effect, which severely compromises the downstream parts of the study. Figure 2a shows examples for such lncRNA-targeting gRNAs, which exhibit off-target activity against well-known housekeeping genes, including RPL3, RPS8, TOP3A, and POLR1. Perturbation of these common essential genes causes severe fitness defects across large cell line panels (Figure 2b, DepMap data), highlighting the magnitude of the confounding effects introduced by such off-targets. The sequence similarity of respective gRNAs to their off-targets is shown in Figure 2c.

**Figure 2:**
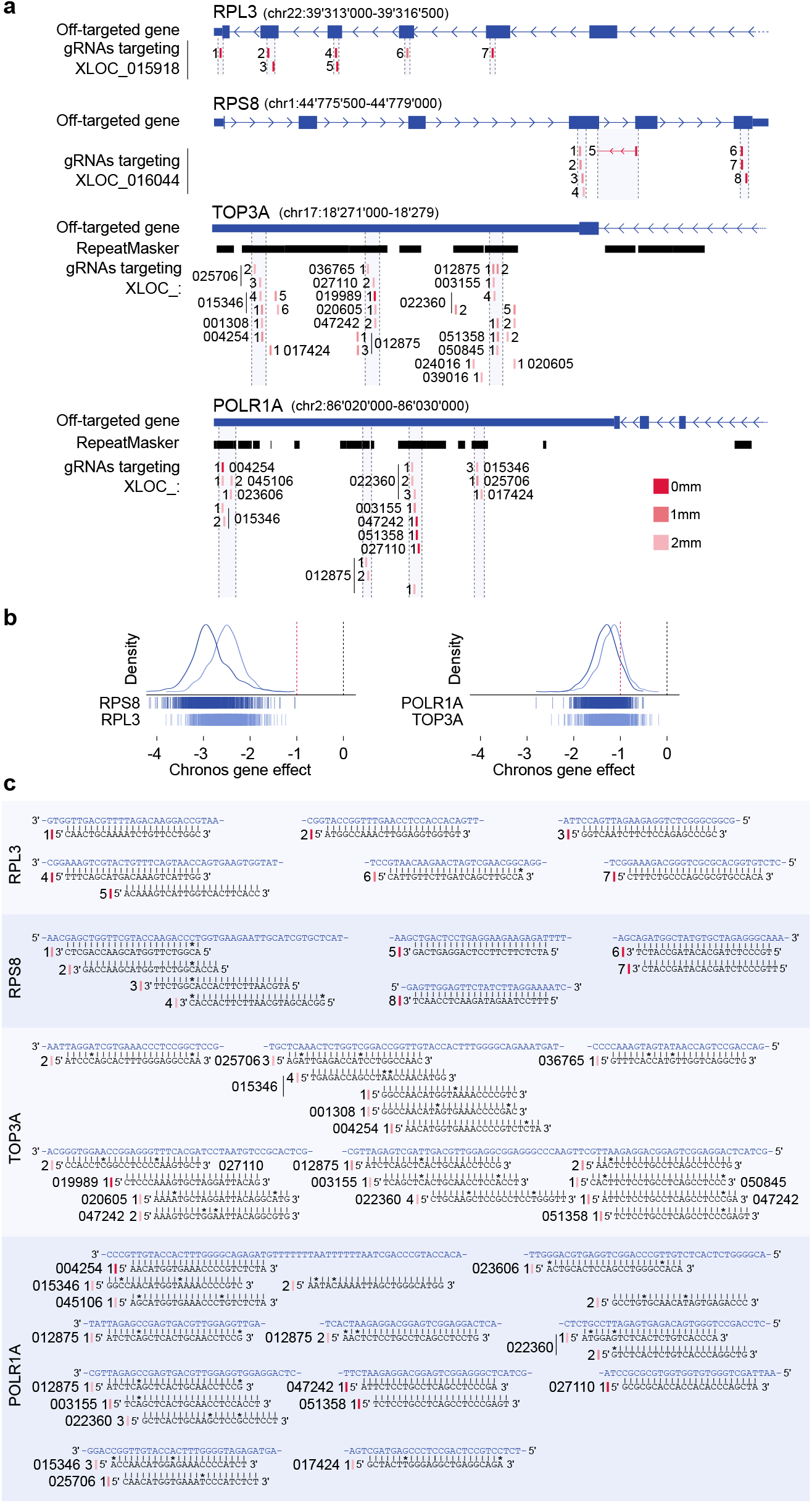
Examples of gRNAs in Liang et al.’s library displaying off-targets in essential protein-coding genes. **a**, Four exemplary essential protein-coding genes affected by off-target gRNAs. The intended lncRNA targets (XLOC_015918 and XLOC_016044) are pseudogenes of *RPL3* and *RPS8*, which encode for ribosomal proteins. *TOP3A* and *POLR1A* are off-targets of gRNAs designed against numerous lncRNAs, as lncRNAs and mRNAs of these protein-coding genes share a SINE repeat in their sequences. The red color scheme indicates gRNAs with a perfect match to their off-targets (0mm) and gRNAs aligning with one (1mm) or two (2mm) mismatches. **b**, DepMap Chronos scores (fitness defect in Cas9 screens) for indicated essential protein-coding genes across all cell lines in DepMap release 24Q2 (n = 1,150 cell lines). **c**, Alignment of gRNA sequences (black) from (a) to the corresponding essential protein-coding gene sequences (blue). Dashes represent perfect matches, asterisks indicate mismatches. Numbering of gRNAs and red color scheme indicating sequence similarity (0mm, 1mm, 2mm) as in (a).

### The majority of “shared lncRNA vulnerabilities” are false-positive

Given that individual lncRNAs are targeted by multiple gRNAs (which might be on- and/or off-target), we next examined how off-target effects impact on the discovery of lncRNA essentialities. To this end, we assigned off-target information to each lncRNA identified as a hit by Liang et al. We found that 55.1% of the 778 lncRNAs identified as hits are targeted by at least one gRNA with an off-target in an essential protein-coding gene (Figure 3a, Table S3). This problem was most pronounced among the “partially shared” and “shared” lncRNA hits, which were affected in 72.5% and 80.4% of cases, respectively (Figure 3a). Notably, these lncRNAs were often targeted by more than one off-target gRNA (Figure 3b).

**Figure 3:**
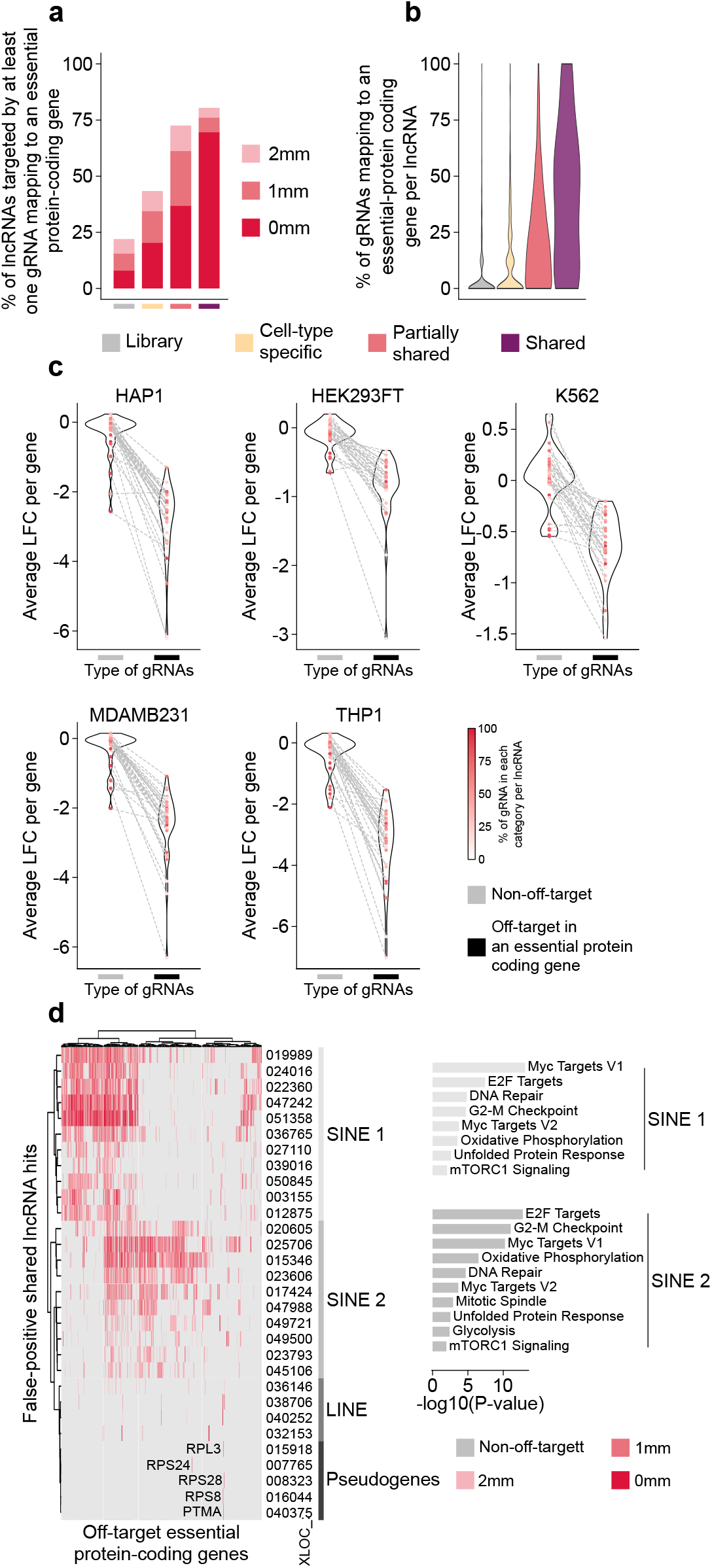
Large fractions of lncRNA hits reported by Liang et al. are false positives due to off-target gRNAs. **a**, Percentage of lncRNAs targeted by at least one gRNA with an off-target effect in an essential protein-coding gene. The information is provided for all lncRNAs in Liang et al.’s library (grey) as well as for the 778 lncRNA hits reported by Liang et al., subdivided by their classification as cell-type-specific (yellow), partially shared (pink), or shared (purple). Red color scheme highlighting the degree of gRNA/off-target sequence similarity (0mm, 1mm, 2mm) as in Figure 1d-f. **b**, Percentage of gRNAs per lncRNA having an off-target in an essential protein-coding gene. For each lncRNA a value was calculated representing the fraction of its gRNAs with off-targets in essential-protein coding genes. Categories and color scheme as in (a). **c**, Cellular fitness defects induced by off-target and non-off-target gRNAs. Data are shown for the 33 “shared” lncRNAs that are targeted by both types of gRNAs. Graphs compare average LFCs caused by off-target gRNAs and non–off-target gRNAs. Dotted lines connect off-target and non-off-target LFC values caused by gRNAs targeting the same lncRNA. These analyses allow classification of “false-positive” hits (off-target gRNAs are the only ones creating fitness defects) or “rescued true-positive” hits (both the off-target and the non-off-target gRNAs caused fitness defects). **d**, Left panel: Heatmap displaying essential protein-coding genes (x-axis) that are off-targets of at least one gRNA targeting “shared” lncRNAs (y-axis). All “shared” lncRNA hits classified as false positives (30/46) are displayed upon applying hierarchical clustering. Red color scheme highlights the degree of gRNA/off-target sequence similarity (0mm, 1mm, 2mm) as in Figure 1d-f. lncRNAs targeted by gRNAs with off-targets in SINE and LINE repeat elements overlapping essential protein-coding genes are indicated. Right panel: Enrichr analyses for off-target essential protein-coding genes falling into indicated groups.

Since off-target effects to essential protein-coding genes are most commonly affecting “shared lncRNA hits”, we examined how related off-target gRNAs influence the fitness phenotypes in this group. We found that among the 46 shared lncRNA hits reported by Liang et al., 4, 9, and 33 were targeted exclusively by off-target gRNAs, exclusively by non-off-target gRNAs, or by both, respectively. Focusing on the latter (lncRNAs targeted by both types of gRNAs), we examined how often the observed fitness defect arose from off-target but not on-target effects (connected by dotted lines, Figure 3c). Indeed, for the majority (81.1%) of these lncRNAs, non-off-target gRNAs showed no detectable phenotype, whereas off-target gRNAs typically produced a strong negative LFC (Figure 3c, Table S3). Thus, fitness defects in these cases are driven exclusively by off-target effects, further supporting the notion that these lncRNAs are false-positive. Only seven lncRNAs exhibited a negative LFC when targeted by both off-target and non-off-target gRNAs (Figure 3c), suggesting that they might be true positives (hereafter referred to as “rescued true-positive hits”). Together, the “rescued true-positive hits” and the hits that are not confounded by off-target effects amount a total of 16 true-positive “shared” lncRNA hits in the screens performed by Liang et al., whereas 30 (65.2%) of the 46 “shared” hits are false-positive. Extension of these analyses to the “partially shared” and “cell type specific” hits revealed that 53.0%, and 33.5%, are false-positives due to off-target effects, respectively (Figures S1a,b, Table S3).

The high frequency of false-positive hits in the screens led us to consider whether there might be common sources for this problem. We therefore examined if there are common features shared by false-positive lncRNAs. Focusing again on the “shared” lncRNA hits, we identified two groups: First, 16.7% of false-positive lncRNAs are in fact pseudogenes with high homology to essential protein-coding genes. Examples for the latter are ribosomal proteins, such as RPL3, RPS24, and RPS28 (Figure 3d, Table S4). Second, 83.3% of false-positive lncRNA genes overlap with repeat elements, including SINE and LINE, which are abundantly present in the genome. Consequently, gRNAs targeting repeat elements have tens to hundreds of off-targets in essential protein coding-genes (Table S4). This explains why some lncRNAs have numerous shared off-target protein-coding genes (Figure 3d, Table S4). Targeting lncRNAs in these two categories is challenging or even impossible in some cases, requiring careful consideration in the future design of Cas13d libraries.

In summary, a large part of lncRNA hits reported by Liang et al. are false-positives due to off-target effects, especially in the groups of “shared” and “partially shared” lncRNA hits. Given that most of the biological findings in the study are based on these groups, the conclusions of the study need to be revisited.

### Validation experiments also suffer from off-target effects

Liang et al conducted a series of experiments to validate their screens (summarized in Figure 4a), which however did not uncover the large number of false-positive hits. Overall, their experiments validated 53 lncRNA hits, primarily “partially shared” or “shared” vulnerabilities, 69.8% of which are clearly false-positive in our analyses. To explore the causes of this discrepancy we examined whether the validation experiments conducted by Liang et al. also suffer from off-target effects. We therefore queried potential off-targets for all gRNAs and DsiRNAs that were used for validation studies, as described above and in the methods section. Overall, we found that 75.5% of lncRNAs studied by Liang et al in their validation experiments were targeted by at least one gRNA or DsiRNA with off-targets in essential protein-coding genes (Figure 4b, Table S5). Exemplary gRNAs and DsiRNAs (designed by Liang et al to target lncRNAs in validation experiments) with off-target activity against common-essential protein-coding genes are shown for the essential-protein coding genes RPS28 and DYNC1H1 in Figure 4c,d. Perturbation of these common essential genes causes severe fitness defects across large cell line panels (DepMap), reinforcing the detrimental nature of such off-target effects.

**Figure 4:**
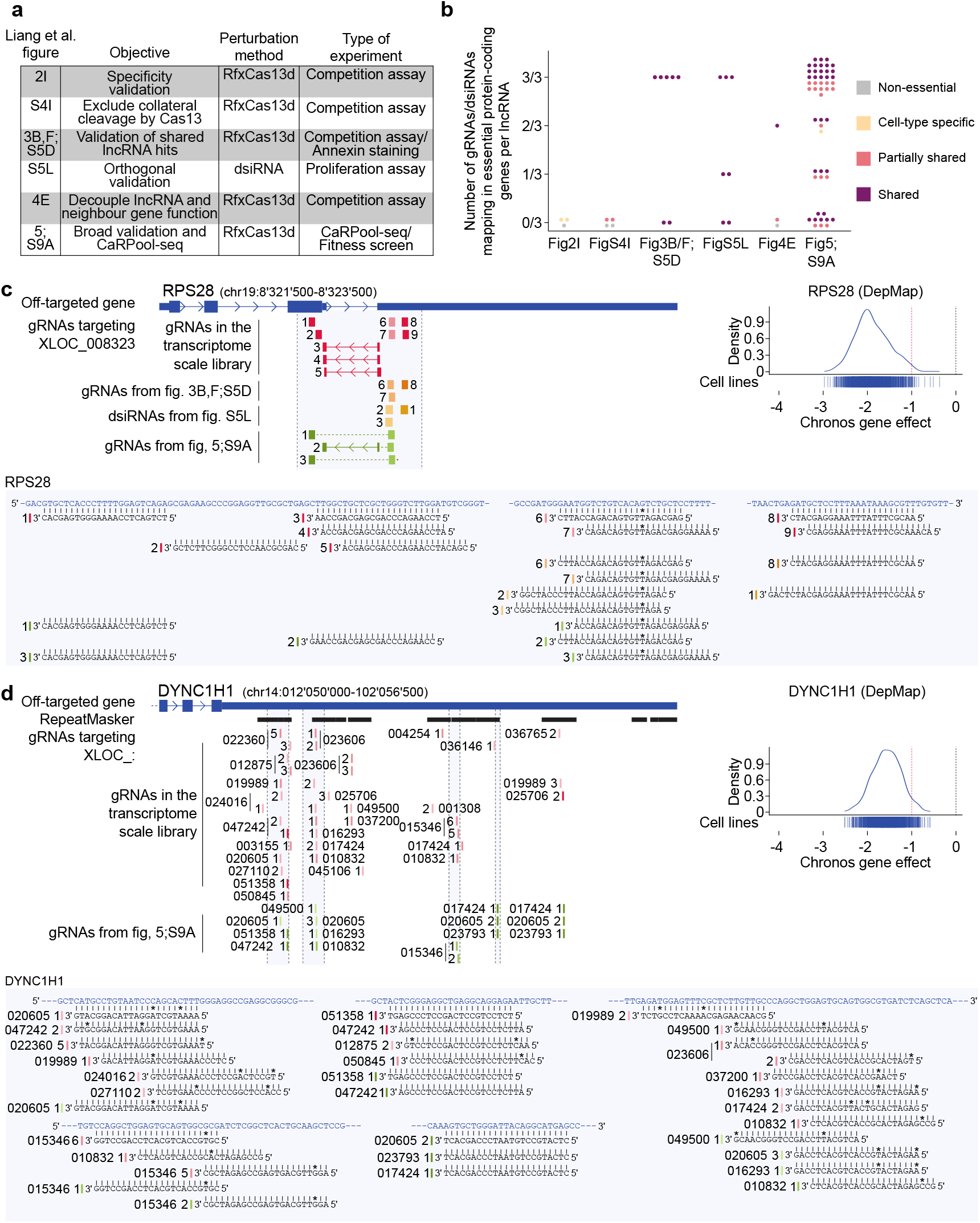
Validation experiments reported by Liang et al. are also compromised by off-target effects. **a**, Overview of validation experiments conducted by Liang et al. **b**. Off-target issues in validation experiments conducted by Liang et al. Liang et al. designed three different gRNAs/DsiRNAs to target each lncRNA. The figure shows for each perturbed lncRNA (represented by dots) the number of gRNAs/DsiRNAs with off-targets in essential protein-coding genes. The color scheme indicates whether individual lncRNAs were initially classified as cell-type-specific (yellow), partially shared (pink), or shared (purple) by Liang et al. **c-d**, Representative examples of essential protein-coding genes targeted by off-target gRNAs/DsiRNAs used in validation experiments by Liang et al. **c**, gRNAs originally designed to target the lncRNA XLOC_008323 have off-targets in *RPS28*. LncRNA XLOC_008323 is a pseudogene with high sequence homology to *RPS28*. **d**, gRNAs designed against multiple lncRNAs have off-targets in *DYNC1H1*, as both the respective lncRNAs and the mRNA of this essential protein-coding gene contain a SINE repeats in their sequences. DepMap Chronos scores indicate fitness defects in Cas9 screens caused by perturbation of *RPS28* and *DYNC1H1* (DepMap release 24Q2, n = 1,150 cell lines).

To illustrate the magnitude of the problem in more detail, we focused on the three validation experiments that encompass the vast majority of studied lncRNAs (Figures 3B/F;S5D, Figure S5L and Figures 5;S9A in Liang et al.). First, experiments shown by Liang et al in their Figures 3B/F and S5D tested 7 “shared” lncRNA hits using Cas13d. We found that three of these lncRNAs were pseudogenes of highly essential protein-coding genes and another two are located in repetitive regions. All gRNAs targeting these lncRNAs in validation experiments showed off-target activity (Figure 4b, Table S5). Only gRNAs against two known cancer-associated lncRNAs, MALAT1 and MIR17HG, did not have off-targets. Second, in the experiments shown by Liang et al in their Figure S5L, the authors validated the same 7 lncRNAs using DsiRNAs as an orthogonal method. However, 5 of those lncRNAs were targeted by at least one DsiRNAs harboring off-targets in essential protein-coding genes (Figure 4b, Table S5), indicating that the conclusions drawn from this orthogonal validation method are also compromised. Third, in their Figures 5 and S9A, Liang et al. present validation experiments for 50 hits - mostly “partially shared” and “shared” lncRNA essentialities. Thereby, they combined Cas13d-based screening with single-cell RNA-Seq using CaRPool-seq, allowing them to study transcriptional changes caused by lncRNA perturbations at the single cell level. We found, however, that, 12%, 10%, and 58% of lncRNAs were targeted by one, two, or three gRNAs with off-target effects in essential protein-coding genes, respectively (Figure 4b, Table S5).

**Figure 5:**
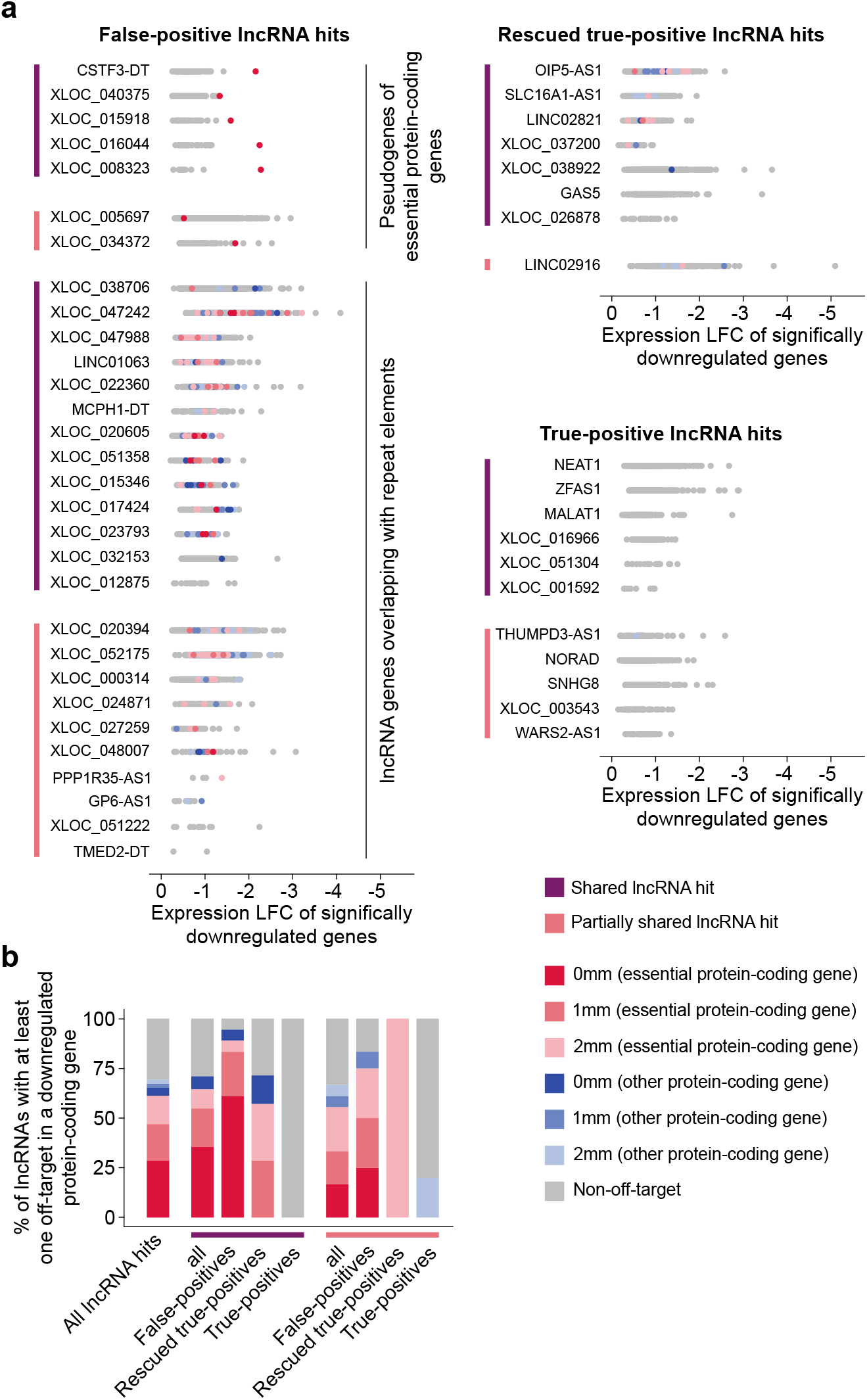
Single-cell transcriptomics combined with Cas13d-mediated lncRNA perturbation confirms downregulation of essential protein-coding genes by off-target gRNAs. Liang et al. performed CaRPool-seq experiments combining single-cell RNA-seq and Cas13d-based lncRNA perturbation. The CaRPool-seq gRNA library was designed to target 50 lncRNAs, which they identified as hits in their primary screens. **a**, List of LncRNAs whose perturbation is associated with significant downregulation of at least one gene (data from CaRPool-seq experiment performed in cell line MDA-MB-231). For each lncRNA perturbation, all significantly downregulated genes (represented by dots) and respective log fold-change values are shown. Colored dots highlight protein-coding genes that are targeted by off-target gRNAs in the CaRPool-seq library. Colors indicate common essential (red) or non-essential protein-coding genes (blue). Different color intensities signify the level of gRNA/off-target similarity (0mm, 1mm, 2mm) as indicated. LncRNAs are assigned to three groups depending on whether they were considered “true-positive”, “rescued true-positive”, or “false-positive” in the evaluation (see Figure 3c and Supplementary Figure 1) of Liang et al.’s primary screens. **b**, Percentage of lncRNAs targeted by at least one gRNA with off-targets in significantly down-regulated protein-coding genes. For each lncRNA the most “impactful” off-target type from (a) was chosen. The strength of impact is indicated by the color gradient (strongest: dark red; weakest: light blue). Values are shown for indicated lncRNA categories.

Overall, these analyses reveal that not only the primary screens but also the validation experiments are severely affected by off-target issues. Less than 25% of validated lncRNAs are bona fide vulnerabilities, while results emanating from the remaining ones need to be revisited.

### CaRPool-seq confirms the downregulation of off-target genes

To evaluate whether gRNAs with off-target effects indeed dysregulate the predicted protein-coding off-target genes in validation experiments, we leveraged Liang et al.’s CaRPool-seq data. Thereby, we focused on the expression of genes that were significantly downregulated following Cas13d-based perturbation (Table S4 from Liang et al). For each lncRNA targeted by the CarPool-seq library, we overlapped the downregulated genes with the predicted off-targets genes (Figure 5a, Table S6).

Overall, these analyses revealed that for 69.4% of lncRNAs the created perturbation downregulated at least one off-target (in most cases essential) protein-coding gene (Figure 5b). This effect was particularly pronounced in the group of lncRNAs defined as false-positive in the primary screens (Figure 5a,b). In this group, downregulation of at least one off-target essential protein-coding gene was observed for 83.3% of lncRNA hits. In contrast, the lncRNA hits classified as “true positives” were not associated with downregulation of off-target essential genes in the CaRPool-seq experiment. This group encompasses mostly lncRNAs that had been previously reported to affect cell proliferation or survival, including NEAT1, ZFAS1, MALAT1, THUMPD3-AS1, NORAD, and SNHG8 (Figure 5a,b). In conclusion, these results confirm that gRNAs with off-target effects cause downregulation of essential protein-coding genes - further supporting the notion (Figure 3) that gRNAs with off-targets are primary drivers of false-positive lncRNA hits.

## Discussion

Systematic mapping of lncRNA vulnerabilities in cancer relies on the application of high-throughput screening approaches. Large-scale targeting of RNA transcripts by Cas13d can overcome some of the limitations of DNA-directed perturbation^9^. However, as with other screening methods, off-target effects can be a confounding factor in Cas13d-based screens. Since gene essentialities are far less common among lncRNAs as compared to protein-coding genes, the relative impact of such confounders can be particularly pronounced in lncRNA screens. This is evident in Liang et al.’s experiments, which suffer considerably from off-target effects in essential protein-coding genes – a problem caused by imperfect gRNA filtering during library design. Consequently, large numbers of reported lncRNA vulnerabilities are false-positive. If left unacknowledged, this problem will negatively impact lncRNA research in different ways.

Considering the limited number of essential lncRNAs described to date, the long list of false-positives among the 778 hits reported by Liang et al. threatens to misdirect lncRNA research for years to come. Overall, 65.2%, 53.0% and 33.5% of the shared, partially shared and cell-type-specific lncRNA hits are caused by off-targets in essential protein-coding genes. Essential lncRNAs are promising targets for drug development in cancer and other diseases, especially since the field of RNA therapeutics is currently undergoing a dynamic phase of expansion and innovation^13^. Pursuing false-positive hits could therefore not only affect basic non-coding RNA research but also misguide translational efforts in biomedicine.

The issues created by off-target effects also compromise the biological conclusions of the study. For example, Liang et al. observed that perturbing essential lncRNAs altered the same proliferation-related pathways. However, this conclusion is likely driven by the similar pattern of off-targets across false-positive hits, causing collateral downregulation of the same essential protein-coding genes. The authors also described that the nearest protein-coding genes of essential lncRNAs are often non-essential - and concluded that these lncRNAs might act independently of nearby genes. Since many “essential” lncRNAs are likely false positives, with fitness effects stemming from distant essential genes, this conclusion should be reassessed. Finally, Liang et al. examined lncRNA expression changes during development and tumorigenesis, concluding that essential lncRNAs are broadly expressed at early developmental stages and are differentially expressed in tumors, correlating with survival. However, these findings strongly relied on comparisons among the expression of shared, partially shared, and cell-type–specific essential lncRNAs, as well as non-essential lncRNAs. Since a large part of hits in all groups are false positive, the conclusions have to be reevaluated upon removal of off-target effects.

In summary, Cas13d based screening^9,10^ will undoubtably advance lncRNA research. Whilst Liang et al’s screening experiments are of high quality, the large number of false-positive lncRNA hits reported in the study undermine the paper’s conclusions and pose a risk to future research building on their results. The reanalysis of screens upon off-target removal will improve the quality of data and provide valuable information to the community.

## Acknowledgements

This study was supported by the European Research Council (Consolidator grant CoG PACA-MET-819642 and MSCA-ITN-ETN-861196 to R.R.; the Deutsche Forschungsgemeinschaft (DFG RA1629/2-1;SFB1321 to R.R.; German Cancer Consortium, Deutsche Krebshilfe (70114314 to R.R.); the German Federal Ministry of Education and Research (Cluster4Future: CNATM to R.R.) and TUM Innovation Network NextGenDrugs funded under the Excellence Strategy of the Federal Government and the Länder (to R.R.). J.J.M. was funded by a European Molecular Biology Organization (EMBO) Long-Term Fellowship (ALFT 655-2019).

## Author Contributions Statement

J.J.M. and R.T. analyzed the data. J.J.M. and R.T. prepared the figures. J.J.M. and R.R. wrote the manuscript.

## Data availability

The sequence of the protein-coding genes RNAs was obtained from GENCODE v38 genome: https://ftp.ebi.ac.uk/pub/databases/gencode/Gencode_human/release_38/gencode.v38.pc_transcripts.fa.gz.

The list of common essential protein-coding genes was obtained from the DepMap portal, release 24Q2: https://depmap.org/portal/data_page/?tab=allData&releasename=DepMap%20Public%2024Q2&filename=CRISPRInferredCommonEssentials.csv.

The coordinates of the lncRNAs targeted by the library of Liang et al. were retrieved from the supplementary data 1 from Sarropoulos et al., 2019^14^:

https://static-content.springer.com/esm/art%3A10.1038%2Fs41586-019-1341-x/MediaObjects/41586_2019_1341_MOESM4_ESM.zip.

---

**Supplementary Figure 1:**
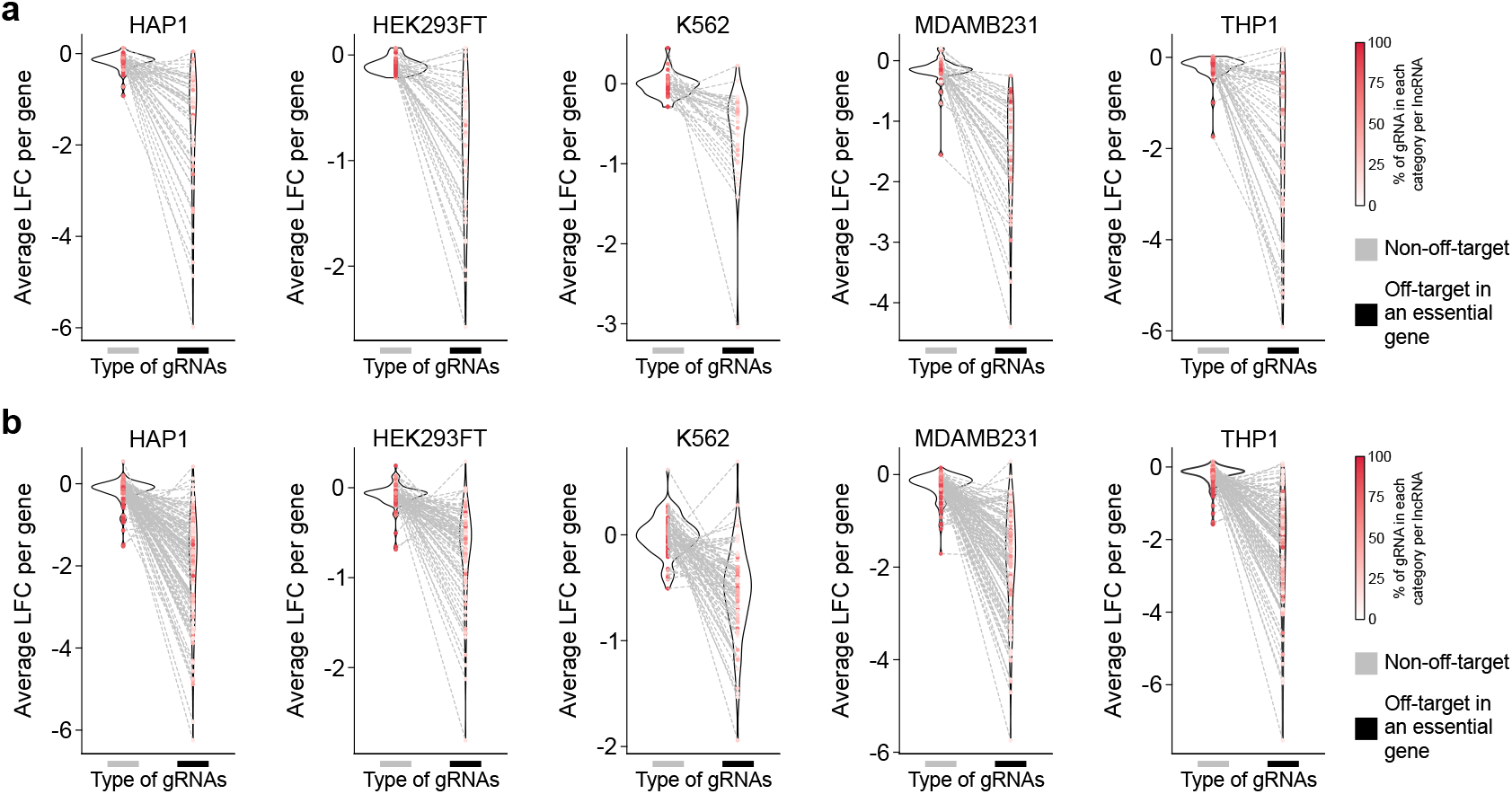
Several partially shared and cell-type-specific lncRNAs are false-positive hits due to off-targets. **a-b**, Comparison of average LFCs caused by off-target gRNAs and non–off-target gRNAs. Only lncRNAs targeted by both on- and off-target gRNAs were considered for these analyses. Off-target and non–off-target average LFC values for the same lncRNA are connected by dashed lines. Data are shown for partially shared (**a**) or cell-type-specific (**b**) lncRNA hits. These analyses allow classification of lncRNAs as “false-positives” (only off-target gRNAs create fitness defects) or “rescued true-positive” (both the off-target and the non–off-target gRNAs cause fitness defects).

## Methods

### Retrieval of lncRNA expression and significance of screening hits from Liang et al

For each lncRNA, expression levels and p-values (indicating significance of enrichment/depletion in the screen) were obtained from Liang et al.’s Tables S1 and S2, respectively. Following the criteria described by Liang et al., a lncRNAs was considered expressed in a given cell line if the corresponding TPM (transcript per million) value was >0, and considered a significant hit in a given screen if the respective p-value was ≤ 0.05 (Table S1).

### Identification of gRNAs with off-target effects

To evaluate if gRNAs from Liang et al. map to protein-coding genes, we first retrieved all mRNA transcript sequences from GENCODE v38^15^. Subsequently, we downloaded the list of common essential protein-coding genes from the DepMap portal (DepMap release 24Q2)^12^. Using this information, we generated two different Bowtie indexes: one using all retrieved protein-coding gene mRNA transcripts, and a second using only protein-coding genes considered common essential in DepMap. Finally, we aligned all lncRNA-targeting gRNA sequences from Liang et al.’s transcriptome-scale Cas13d CRISPR library (Table S1 from Liang et al.^10^) to the complementary strand (--nofw) of the mRNA transcriptome references using Bowtie (v1.2.2) (https://doi.org/10.1186/gb-2009-10-3-r25), allowing up to two mismatches (-v 2):

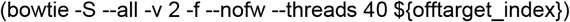

A gRNA was considered to suffer from an off-target effect against a particular protein-coding gene if it mapped (Bowtie) to at least one of its transcript isoforms using the parameters described above. We then retrieved the “quality of the mapping” (number of mismatches) and determined whether the off-target gene was an essential protein-coding gene or not (Table S2). The same procedure was applied to Cas13d gRNAs used in the validation experiments from Liang et al. (Table S5).

### Identification of DsiRNAs with off-target effects

To evaluate the potential off-target effects of DsiRNAs used in the orthogonal validation experiment performed by Liang et al., we used BLAST (v2.16.0)^16^. We aligned the bottom strand of each 27-mer DsiRNA to the transcriptome of all the protein-coding genes or to essential protein-coding genes (using the same mRNA sequences described above) using blastn-short (no DUST filtering and no soft masking). Only alignments on the complementary strand of the mRNAs were taken into consideration. A gRNA was considered to exhibit off-target effects against a protein-coding gene if its sequence mapped to at least one transcript isoform with up to four mismatches and no gaps (Table S5). This more permissive approach was used for DsiRNAs because they are longer than the Cas13d gRNAs and have a shorter seed region (2-7 nucleotides^17–20^) compared to Cas13d gRNAs (the optimum gRNA length is 20–23 nucleotides^21,22^).

### Evaluation of the sequence similarity between lncRNAs and essential protein-coding genes

The sequences of all lncRNAs included in the transcriptome-scale Cas13d CRISPR library from Liang et al. were obtained as follows: First, the genomic coordinates for each lncRNA isoform were obtained from Sarropoulos et al.^14^. Second, these coordinates were used to retrieve the transcript sequence for each lncRNA gene using the getSeq function from the BSgenome.Hsapiens.UCSC.hg19 package (R 4.4.1). With this data, we calculated the sequence similarity between the lncRNAs and all essential protein-coding transcripts. For this, all lncRNA isoform sequences were aligned to the essential transcriptome reference using BLAST^23^ (v2.16.0) with default parameters, excluding alignments on the reverse strand. When multiple isoforms of the same lncRNA yielded different similarity scores, the highest value was retained for downstream analyses (Table S4).

### Gene set enrichment analysis

All essential protein-coding genes affected by at least one off-target gRNA designed against a lncRNA included in the SINE1 and SINE2 clusters (Figure 3d) were retrieved. To reduce noise, only protein-coding genes affected by off-target gRNAs designed to perturb at least two different lncRNAs were retained. Gene set enrichment analysis was then performed using Enrichr^24^, and results were retrieved only for the Hallmark gene sets, as these were the pathways used by Liang et al. to construct their Figure 5D. Only gene sets with an FDR-adjusted p-value < 0.05 were considered for further visualization (Figure 3d).

## References

1. Long non-coding RNAs: definitions, functions, challenges and recommendations | Nature Reviews Molecular Cell Biology. https://www.nature.com/articles/s41580-022-00566-8.

2. Ferrer, J. & Dimitrova, N. Transcription regulation by long non-coding RNAs: mechanisms and disease relevance. Nat Rev Mol Cell Biol 25, 396–415 (2024).

3. Ma, L. et al. LncBook: a curated knowledgebase of human long non-coding RNAs. Nucleic Acids Res 47, 2699 (2019).

4. Zhu, S. et al. Genome-scale deletion screening of human long non-coding RNAs using a paired-guide RNA CRISPR-Cas9 library. Nat Biotechnol 34, 1279–1286 (2016).

5. Liu, Y. et al. Genome-wide screening for functional long noncoding RNAs in human cells by Cas9 targeting of splice sites. Nat Biotechnol 36, 1203–1210 (2018).

6. Liu, S. J. et al. CRISPRi-based genome-scale identification of functional long noncoding RNA loci in human cells. Science 355, aah7111 (2017).

7. Boutros, M. & Ahringer, J. The art and design of genetic screens: RNA interference. Nat Rev Genet 9, 554–566 (2008).

8. Lennox, K. A. & Behlke, M. A. Cellular localization of long non-coding RNAs affects silencing by RNAi more than by antisense oligonucleotides. Nucleic Acids Res 44, 863–877 (2016).

9. Montero, J. J. et al. Genome-scale pan-cancer interrogation of lncRNA dependencies using CasRx. Nat Methods 21, 584–596 (2024).

10. Liang, W.-W. et al. Transcriptome-scale RNA-targeting CRISPR screens reveal essential lncRNAs in human cells. Cell 187, 7637-7654.e29 (2024).

11. Guo, X. et al. Transcriptome-wide Cas13 guide RNA design for model organisms and viral RNA pathogens. Cell Genomics 1, (2021).

12. Tsherniak, A. et al. Defining a Cancer Dependency Map. Cell 170, 564-576.e16 (2017).

13. Winkle, M., El-Daly, S. M., Fabbri, M. & Calin, G. A. Noncoding RNA therapeutics — challenges and potential solutions. Nat Rev Drug Discov 20, 629–651 (2021).

14. Sarropoulos, I., Marin, R., Cardoso-Moreira, M. & Kaessmann, H. Developmental dynamics of lncRNAs across mammalian organs and species. Nature 571, 510–514 (2019).

15. Frankish, A. et al. GENCODE 2021. Nucleic Acids Research 49, D916–D923 (2021).

16. Altschul, S. F., Gish, W., Miller, W., Myers, E. W. & Lipman, D. J. Basic local alignment search tool. Journal of Molecular Biology 215, 403–410 (1990).

17. Valencia-Sanchez, M. A., Liu, J., Hannon, G. J. & Parker, R. Control of translation and mRNA degradation by miRNAs and siRNAs. Genes Dev 20, 515–524 (2006).

18. Birmingham, A. et al. 3’ UTR seed matches, but not overall identity, are associated with RNAi off-targets. Nat Methods 3, 199–204 (2006).

19. Jackson, A. L. et al. Widespread siRNA ‘off-target’ transcript silencing mediated by seed region sequence complementarity. RNA 12, 1179–1187 (2006).

20. Lim, L. P. et al. Microarray analysis shows that some microRNAs downregulate large numbers of target mRNAs. Nature 433, 769–773 (2005).

21. Konermann, S. et al. Transcriptome Engineering with RNA-Targeting Type VI-D CRISPR Effectors. Cell 173, 665-676.e14 (2018).

22. Li, S. et al. Screening for functional circular RNAs using the CRISPR-Cas13 system. Nat Methods 18, 51–59 (2021).

23. Altschul, S. F., Gish, W., Miller, W., Myers, E. W. & Lipman, D. J. Basic local alignment search tool. J Mol Biol 215, 403–410 (1990).

24. Kuleshov, M. V. et al. Enrichr: a comprehensive gene set enrichment analysis web server 2016 update. Nucleic Acids Res 44, W90–97 (2016).

