## Supplementary tables 1-6 for "Off-target effects as confounders of Cas13d-based lncRNA screens": Read me.txt

Table S1: List of all lncRNAs included in the transcriptome-scale Cas13d library from Liang et al. The table includes p-values from fitness screens across all cell lines studied by Liang et al., and indicates whether each lncRNA was considered a hit according to the authors� criteria. It also includes the expression level (TPM) for each lncRNA, whether it was classified as "expressed" by the authors, and the original classification of each lncRNA hit based on its recurrence across screens. Additionally, a re-classification is provided using the same criteria, but accounting for cases where a lncRNA was a hit in a cell line where it was not expressed.

Table S2: List of all gRNAs targeting lncRNAs included in the transcriptome-scale Cas13d library from Liang et al. The table includes log-fold change (LFC) values from the screens for each gRNA, whether the gRNA was considered phenotypic, whether it maps to the sequence of protein-coding genes, the IDs of those protein-coding genes, mapping quality, and whether any off-targeted protein-coding genes are essential.

Table S3: Summary of off-targets associated with lncRNAs targeted by gRNAs in the transcriptome-scale Cas13d library from Liang et al. The table lists all protein-coding genes off-targeted by at least one gRNA per lncRNA, mapping quality of these gRNAs to off-target genes, and whether each off-targeted gene is considered a common essential gene. It also compares the average LFC values of off-target versus non�off-target gRNAs for all lncRNA hits targeted by both types of gRNAs.

Table S4: Characterization of the shared lncRNA hits reported by Liang et al. The figure shows the number of common essential protein-coding genes off-targeted by at least one gRNA designed against each lncRNA, the highest sequence similarity of each lncRNA transcript to essential protein-coding mRNAs, and whether the lncRNA is considered a pseudogene of a protein-coding gene or contains a repeat element in its sequence.

Table S5: List of all gRNAs/DsiRNAs used by Liang et al. in their validation experiments. The table indicates whether each gRNA/DsiRNA has an off-target in a protein-coding gene, the number of mismatches, and whether the off-targeted gene is essential.

Table S6: Integration of CaRPool-seq data from Liang et al. in MDA-MB-231 cells with off-target effects in protein-coding genes. The table summarizes the off-targets of gRNAs used in the CaRPool-seq experiment and their impact.
